# ALIX and ceramide differentially control polarized exosome release from epithelial cells

**DOI:** 10.1101/2020.08.05.238931

**Authors:** Takahide Matsui, Shu Hiragi, Futaba Osaki, Yuriko Sakamaki, Mitsunori Fukuda

## Abstract

Exosomes, new players in cell-cell communication, are extracellular vesicles of endocytic origin. Although single cells are known to release various kinds of exosomes (referred to as exosomal heterogeneity), very little is known about the mechanisms by which they are produced and released. Here, we established methods for studying exosomal heterogeneity by using polarized epithelial cells and showed that distinct types of exosomes are differentially secreted from the apical and basolateral sides. We also identified GPRC5C (G protein-coupled receptor class C group 5 member C) as an apical-exosome-specific protein. We further demonstrated that basolateral exosome release depends on ceramide, whereas ALIX, an ESCRT (endosomal sorting complexes required for transport)-related protein, not the ESCRT machinery itself, is required for apical exosome secretion. Thus, two independent machineries, the ALIX–Syntenin1– Syndecan1 machinery (apical side) and the sphingomyelinase-dependent ceramide production machinery (basolateral side), are likely to be responsible for the polarized exosome release from epithelial cells.

## Introduction

Cells release extracellular vesicles (EVs) of different sizes and intracellular origin. Exosomes (around 100 nm in diameter) are EVs derived from multivesicular bodies (MVBs), and they are released by well-organized systems. Exosomal cargos, such as proteins, lipids, and nucleic acids, are selectively incorporated into intraluminal vesicles (ILVs), i.e., precursors of exosomes, in MVBs. The MVBs are then transported to the plasma membrane, and after fusing with it, the ILVs are released into the extracellular space as exosomes (Pegtel & Gould, 2019; Kalluri & LeBleu, 2020). Exosomal heterogeneity, that is, various types (e.g., sizes and contents) of exosomes being released from a single cell, has recently been reported (Colombo *et al*, 2013; Kowal *et al*, 2016; Zhang *et al*, 2018). Although several distinct mechanisms of exosome biogenesis have been reported (Kalluri & LeBleu, 2020; Mathieu *et al*, 2019), how these mechanisms are differently used or regulated within a single cell remains completely unknown. This is mainly because the results of the studies varied with the techniques and devices used to conduct them. Hence, the mechanisms by which heterogenous exosomes are produced within cells is poorly understood.

The MDCK (Madin-Darby canine kidney) cell line is a well-known epithelial cell line, which has clearly defined apical–basolateral asymmetry (i.e., apical and basolateral domains), and for that reason MDCK cells are often used as an *in vitro* model for studying the mechanism of polarization (Simmons, 1982). Once non-polarized cells release heterogenous exosomes into the extracellular space, it is extremely difficult to distinguish and collect them separately. However, if heterogenous exosomes are asymmetrically released from polarized MDCK cells, it would be possible to easily collect apical and basolateral exosomes separately. Thus, we assumed that MDCK cells would become a good model for studying exosomal heterogeneity without using special techniques and devices. Here, we established a method for purifying exosomes released from polarized MDCK cells and found that the polarized cells release distinct types of exosomes (apical and basolateral exosomes) with different protein composition. Moreover, we showed that the ESCRT (endosomal sorting complexes required for transport) machinery is not unexpectedly required for exosome release from polarized MDCK cells, and instead, ALIX and ceramide independently regulate apical and basolateral exosome release, respectively.

## Results and discussion

### Polarized MDCK cells release distinct types of exosomes from apical and basolateral side

To investigate differences between apical and basolateral exosomes, we first purified all EVs from apical and basolateral MDCK culture media by PEG (polyethylene glycol) precipitation (Rider *et al*, 2016; Cocozza *et al*, 2020) (Fig EV1) and found that four well-known exosome marker proteins (Flotillin-1, CD63, CD9, and CD81) were asymmetrically recovered in the apical and basolateral PEG pellets (Fig 1A), consistent with the previous reports (Banfer *et al*, 2018; Chen *et al*, 2016). Moreover, by using a density gradient floatation assay, each exosome marker in the PEG pellets was floated into the same fraction (Fr. 5 in Fig 1B), suggesting that the PEG pellets contained membranous organelles having the same density. Furthermore, an immunofluorescence analysis showed that CD63 (enriched in apical EVs) was not completely overlapped with CD9 (enriched in basolateral EVs) in polarized MDCK cells (Fig EV2A). Taken together, these results allowed us to conclude that a single MDCK cell secretes at least two types of EVs presumably from different origins that have different exosome markers: Flotillin-1- and CD63-enriched vesicles from the apical side, and CD9- and CD81-enriched vesicles from the basolateral side.

**Figure 1.**
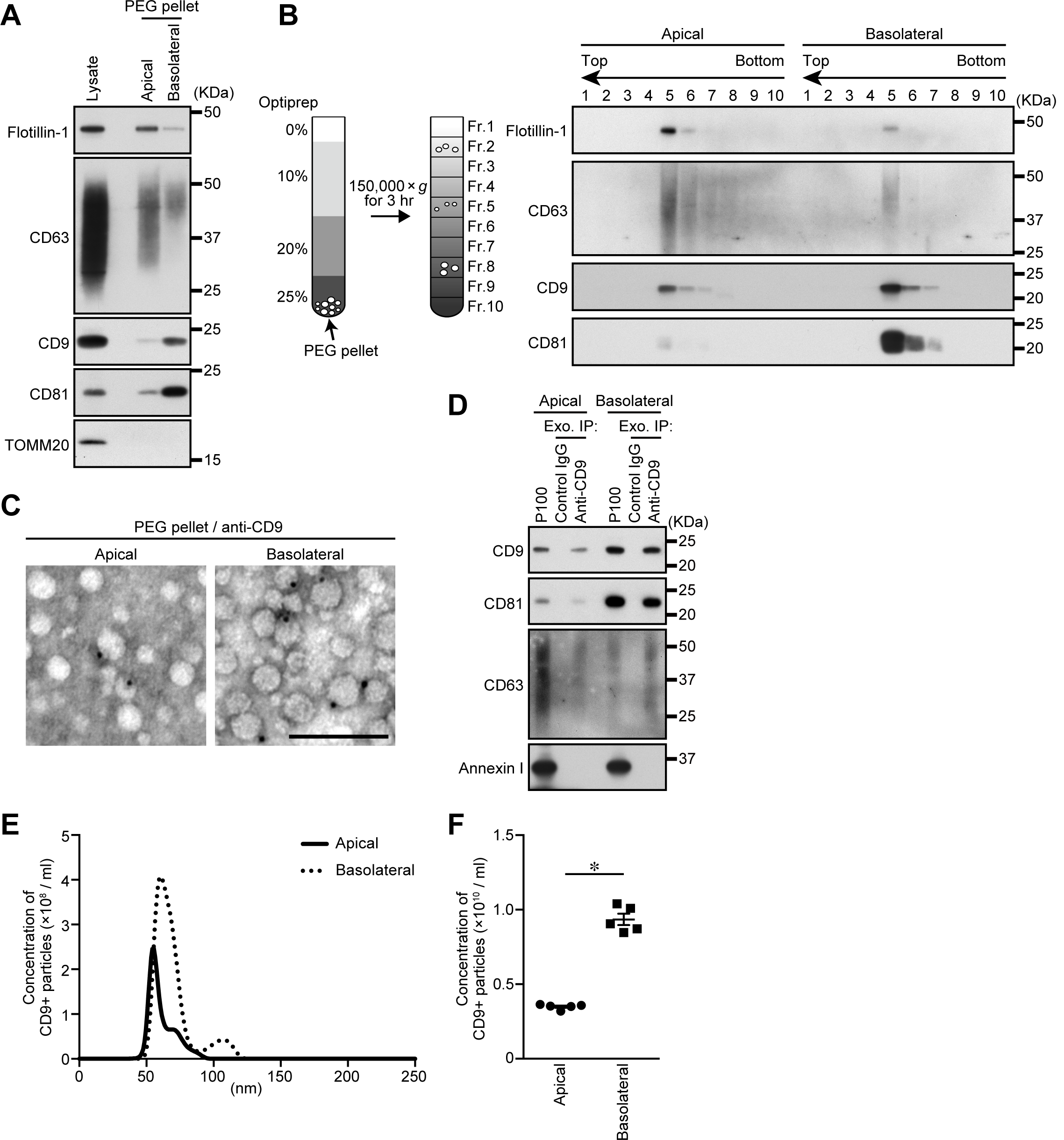
Heterogenous exosome release from polarized MDCK cells. A. MDCK cells were cultured on cell culture inserts for 4 days. On the last day, the culture medium was replaced with EV-depleted medium. EVs released from the apical and basolateral sides of MDCK cells were purified by PEG precipitation. Cell lysates and EV proteins in PEG pellets were analyzed by immunoblotting with the antibodies indicated. Note that the PEG pellets did not contain mitochondrial protein TOMM20, suggesting that the PEG pellets were not contaminated by intracellular organelles. B. PEG pellets prepared as in (A) were subjected to OptiPrep flotation analysis. C. PEG pellets prepared as in (A) were immunonegative stained with anti-CD9 antibody and analyzed by electron microscopy. Scale bar, 100 nm. D. Exosomes were isolated from the pre-cleared medium by direct immunoaffinity capture using anti-CD9 antibody. E. Exosomes prepared as in (D) were eluted from the beads with a glycine buffer and analyzed by nanoparticle tracking analysis (NTA). Representative NTA traces were shown. F. Quantification of the NTA data obtained in five independent experiments. The statistical analyses were performed by two-sided Student’s unpaired *t* test. \**P* < 0.01. Mean ± s.e.m. was shown.

To determine whether these exosome-marker-positive EVs in the PEG pellets are actually exosomes, we initially focused on their size. An electron-microscopic analysis of the MDCK cells revealed that most of the ILVs in their MVBs were less than 100 nm in diameter (Fig EV2B). A nanoparticle tracking assay (NTA), however, showed that both the apical and basolateral PEG pellets contained larger particles as well as 50-100 nm particles, although their size distributions and concentrations were similar (Figs EV2C and D). Furthermore, immunonegative staining of the PEG pellets with anti-CD9 antibody showed that, despite being of similar size, some of the vesicles were CD9-positive, and others were CD9-negative (Fig 1C). These observations indicated that the PEG pellets contained various types of small EVs besides exosomes and thus that vesicle size is inappropriate as a criterion for categorizing EVs, at least exosomes.

To directly analyze exosomes, we next performed direct immunoaffinity capture of exosomes by using anti-CD9 antibody (see (Jeppesen *et al*, 2019) for details) (Fig EV1). As shown in Fig 1D, both apical and basolateral CD9-positive exosomes contain CD81 and CD63, but not Annexin I, a specific marker for microvesicles (Jeppesen *et al*., 2019), and CD9 and CD81 were more abundant in the basolateral exosomes than in the apical exosomes, the same as in the PEG pellets (Fig 1A). Moreover, NTA showed that most of the apical and basolateral CD9-positive exosomes were less than 100 nm in diameter (Fig 1E), consistent with the results of the electron-microscopic analyses (Figs 1C and EV2B), and the concentration of the basolateral CD9-positive exosomes was higher than that of the apical exosomes (Fig 1F). These results suggested that MDCK cells differentially release CD9-positive exosomes from apical and basolateral sides.

### GPRC5C is a novel apical-exosome-specific protein

To further clarify the difference between the apical and basolateral exosomes, we searched for apical- or basolateral-exosome-specific proteins by liquid chromatography-tandem mass spectrometry (LC-MS/MS) (Datasets EV1-3). We used all EVs collected by ultracentrifugation (Fig EV1) to perform LC-MS/MS, because there was too little CD9-positive-exosomes protein to analyze. Silver staining of the proteins from apical and basolateral EVs yielded similar band patterns (Fig 2A), and 84% of the proteins detected by LC-MS/MS in the two types of EVs were identical (Fig 2B). One of the proteins detected, GPRC5C (G protein-coupled receptor class C group 5 member C), was detected only in the apical sample. GPRC5C is an orphan receptor that belongs to the GPRC5 family (Robbins *et al*, 2000) and is involved in renal acid-base homeostasis (Rajkumar *et al*, 2018). but it had never been reported as an exosomal protein. GPRC5C was detected in the apical PEG pellet alone (Fig 2C) and floated into Fr. 5, the same as other exosomal proteins (Fig 2D). Moreover, GPRC5C was co-immunopurified only with apical CD9-positive exosomes (Fig 2E), indicating that GPRC5C is a novel exosomal protein, and that MDCK cells release at least two types of CD9-positive exosomes (a GPRC5C-positive type from the apical side and a GPRC5C-negative type from the basolateral side).

**Figure 2.**
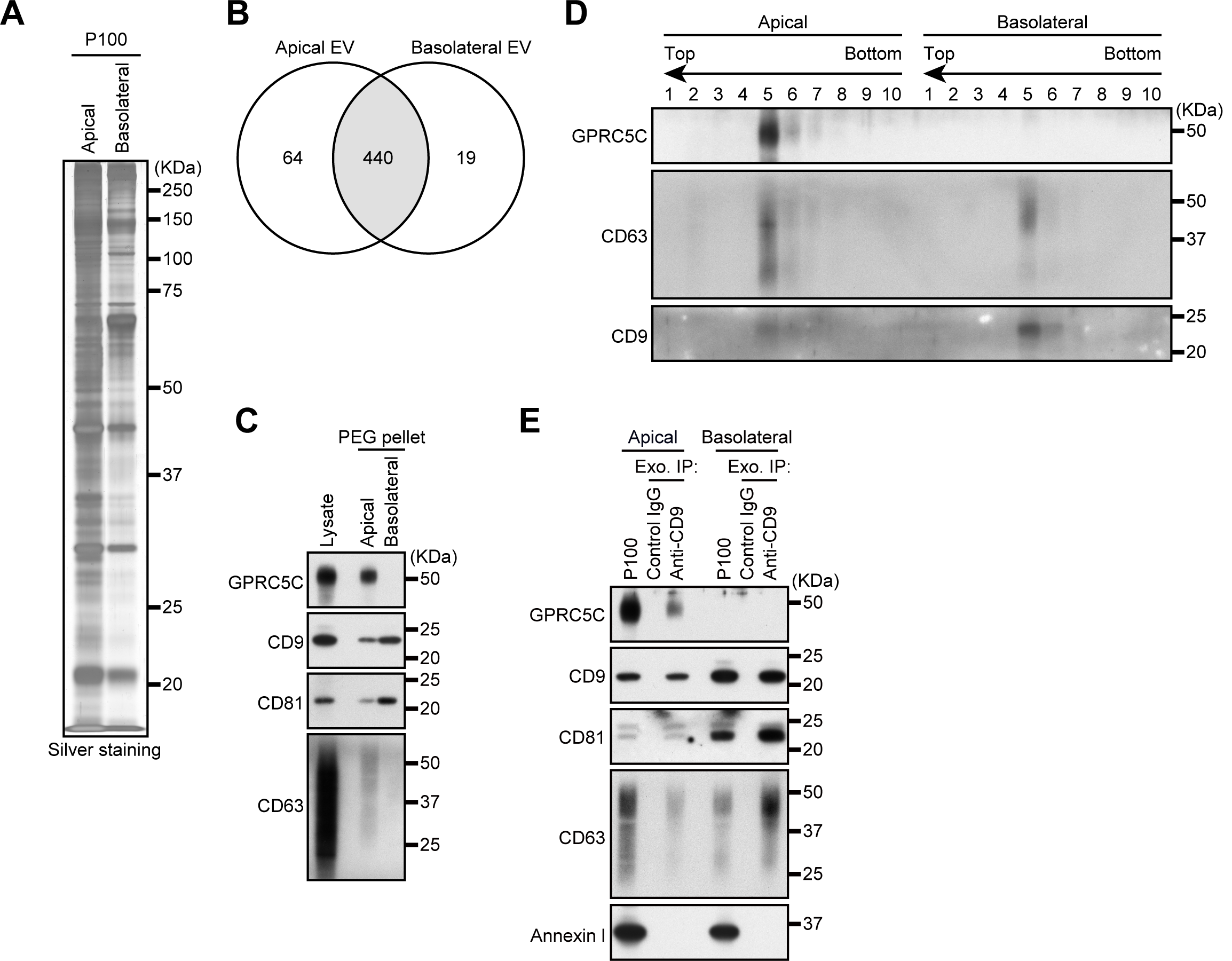
GPRC5C is an apical specific exosomal protein. A. MDCK cells were cultured on cell culture inserts for 4 days. On the last day, the culture medium was replaced with serum-free medium. EVs released from the apical and basolateral sides of MDCK cells were purified by ultracentrifugation. Apical and basolateral EVs (P100) were analyzed by silver staining. B. P100 prepared as in (A) were analyzed by LC-MS/MS. Venn diagrams represent the number of proteins detected in each sample with minimum three peptides. See also Datasets S1-S3. C. MDCK cells were cultured on cell culture inserts for 4 days. On the last day, the culture medium was replaced with EV-depleted medium. EVs released from the apical and basolateral sides of MDCK cells were purified by PEG precipitation. Cell lysates and EV proteins in PEG pellets were analyzed by immunoblotting with the antibodies indicated. D. PEG pellets were subjected to OptiPrep flotation analysis. E. Exosomes were isolated from the pre-cleared medium by direct immunoaffinity capture using anti-CD9 antibody. Note that GPRC5C was detected only in the apical EVs or exosome samples in all experiments performed.

### Inhibition of the ESCRT machinery promotes exosome release

The release of heterogenous exosomes from a single cell requires the production of ILVs or MVBs having different properties. It is well known that the ESCRT machinery regulates ILV formation and cargo sorting into ILVs (Christ *et al*, 2017; Henne *et al*, 2011; Raiborg & Stenmark, 2009; Vietri *et al*, 2020). However, involvement of ESCRT in exosome biogenesis has been controversial, because some groups have reported that MVBs can be generated and exosomes are released in an ESCRT-independent manner (Colombo *et al*., 2013; Trajkovic *et al*, 2008). We therefore attempted to determine whether the ESCRT machinery is involved in the polarized exosome release from MDCK cells. The ESCRT machinery and its associated proteins can be divided into six functionally distinct subcomplexes: ESCRT-0/I/II/III, VPS4, and other ESCRT-associated proteins. We first knocked down the component(s) (HRS (ESCRT-0), TSG101 (ESCRT-I), EAP20 and 30 (ESCRT-II), CHMP6 (ESCRT-III), VPS4A/B, and ALIX (other ESCRT-associated proteins)) of each subcomplex and examined the amounts of exosome markers released (Fig 3A). Knockdown (KD) of most of the ESCRT proteins (HRS, TSG101, EAP20/30, CHMP6, and VPS4A/B) promoted CD9-positive exosome secretion from both sides. Moreover, the numbers of exosomes released from these cells increased without any change in their size (Figs 3B and C). It is generally thought that MVBs fuse with lysosomes for their degradation rather than with the plasma membrane to release exosomes and that they are essential for the lysosomal function via the endocytic pathway (Huotari & Helenius, 2011). Since lysosomal dysfunction has been shown to result in larger MVBs and to promote exosome release from breast cancer cells (Latifkar *et al*, 2019), we hypothesized that exosome release is accelerated by the lysosomal dysfunction caused by ESCRT-KD. To test this hypothesis, we exposed MDCK cells to the vacuolar ATPase inhibitor bafilomycin A_1_ to abolish the lysosomal function and confirmed that CD9-positive exosome release from both sides was upregulated in an exposure-time-dependent manner (Figs EV3A-C). Moreover, enlarged MVBs were observed in the bafilomycin A_1_-exposed cells, the same as in the HRS-KD and VPS4-KD cells (Figs EV3D and E) (Stuffers *et al*, 2009), indicating that most of the ESCRTs-KD affect the lysosomal function, thereby promoting exosome release.

**Figure 3.**
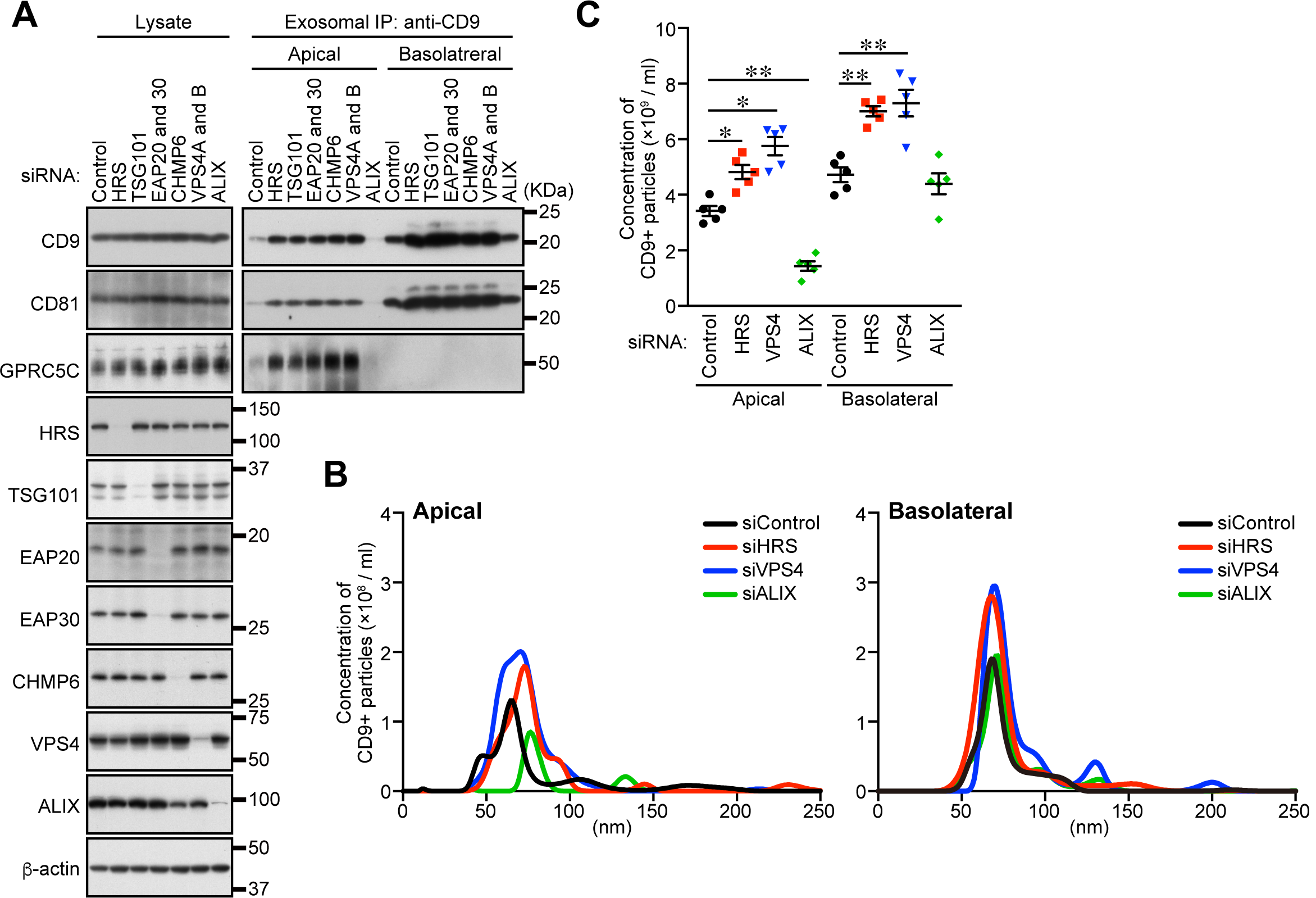
ALIX, but not the ESCRT machinery, is required for apical exosome release. A. MDCK cells were transfected with siControl or the siRNAs indicated, and the cells were transferred to cell culture inserts and cultured for 4 days. On the last day, the culture medium was replaced with EV-depleted medium. Exosomes were isolated from the pre-cleared medium by direct immunoaffinity capture using anti-CD9 antibody. Cell lysates and exosome samples were analyzed by immunoblotting with the antibodies indicated. B. Exosomes prepared as in (A) were eluted from the beads with a glycine buffer and analyzed by NTA. Representative NTA traces were shown. C. Quantification of the NTA data obtained in five independent experiments. \**P* < 0.05, \*\**P* < 0.01 (one-way ANOVA and Tukey’s test). Mean ± s.e.m. was shown.

### Depletion of ALIX specifically reduces apical exosome release

Unlike these ESCRT-KDs, ALIX-KD specifically decreased apical exosome release, and did not affect basolateral exosome release (Figs 3A-C and EV3G). Because the size of the MVBs in the ALIX-KD cells seemed to be unaffected, unlike the HRS-KD or VPS4-KD cells (Figs EV3E and F), ALIX is likely to be involved in apical exosome release independently of the ESCRT machinery. Actually, ALIX is also known to form a ternary complex with Syntenin1 and Syndecan1 and to regulate exosome biogenesis (Baietti *et al*, 2012; Ghossoub *et al*, 2014). As shown in Figure S4, Syntenin1-KD and Syndecan1-KD phenocopied ALIX-KD (Figs 3A-C and EV3G), strongly suggesting that ALIX regulates apical exosome release together with Syntenin1 and Syndecan1, and independently of the ESCRT machinery. Since ALIX has been shown to mediate the sorting of exosomal cargo proteins to ILVs (Dores *et al*, 2012; Dores *et al*, 2016; Larios *et al*, 2020) and Syntenin1 can bind CD63 (Latysheva *et al*, 2006), an abundant protein in apical exosomes (Fig. 1), the ALIX–Syntenin1–Syndecan1 complex presumably regulates the cargo protein sorting to apical exosomes and may also be involved in the efficient ILV formation in MDCK cells.

### Inhibition of ceramide synthesis specifically inhibits basolateral exosome release

To identify the mechanism of the ALIX/ESCRT-independent basolateral exosome release, we turned our attention to the sphingolipid ceramide, because it is enriched in exosomes and regulates ILV formation, cargo sorting, and exosome release independently of the ESCRT machinery (Trajkovic *et al*., 2008). Ceramide is formed as a result of the hydrolytic removal of the phosphocholine moiety of sphingomyelin by sphingomyelinases (SMases), and the neutral SMase inhibitor GW4869 is often used as an effective drug to suppress exosome release (Trajkovic *et al*., 2008; Catalano & O’Driscoll, 2020; Verweij *et al*, 2018). When MDCK cells were treated with GW4869, basolateral CD9-positive exosome release was specifically reduced without affecting apical CD9-positive exosome release (Figs EV5A-C). Essentially, the same results were obtained by nSMase2-KD (Figs EV5D-F), indicating that ceramide is involved only in basolateral exosome release.

### ALIX and ceramide mediate polarized exosome release independently of each other

Finally, we investigated whether ALIX and ceramide mediate the polarized release of exosomes from MDCK cells independently of each other. The results showed that ALIX-KD reduced the amount of the apical exosome marker and number of exosomes released, and that GW4869 had no effect, whereas the opposite results were obtained regarding basolateral exosomes (Figs 4A-C). Since no accumulation of exosome markers in total lysates was observed even in ALIX-KD and GW4869-treated cells, secretory MVBs may represent only a minor population of all MVBs. Alternatively, inhibition of exosome release may promote degradation of secretory MVBs, in contrast to lysosomal dysfunction, which promoted exosome release (Figs EV3A-C). Taken together, these results indicated that ALIX and ceramide independently mediate apical exosome release and basolateral exosome release, respectively (Fig 4D).

**Fig. 4.**
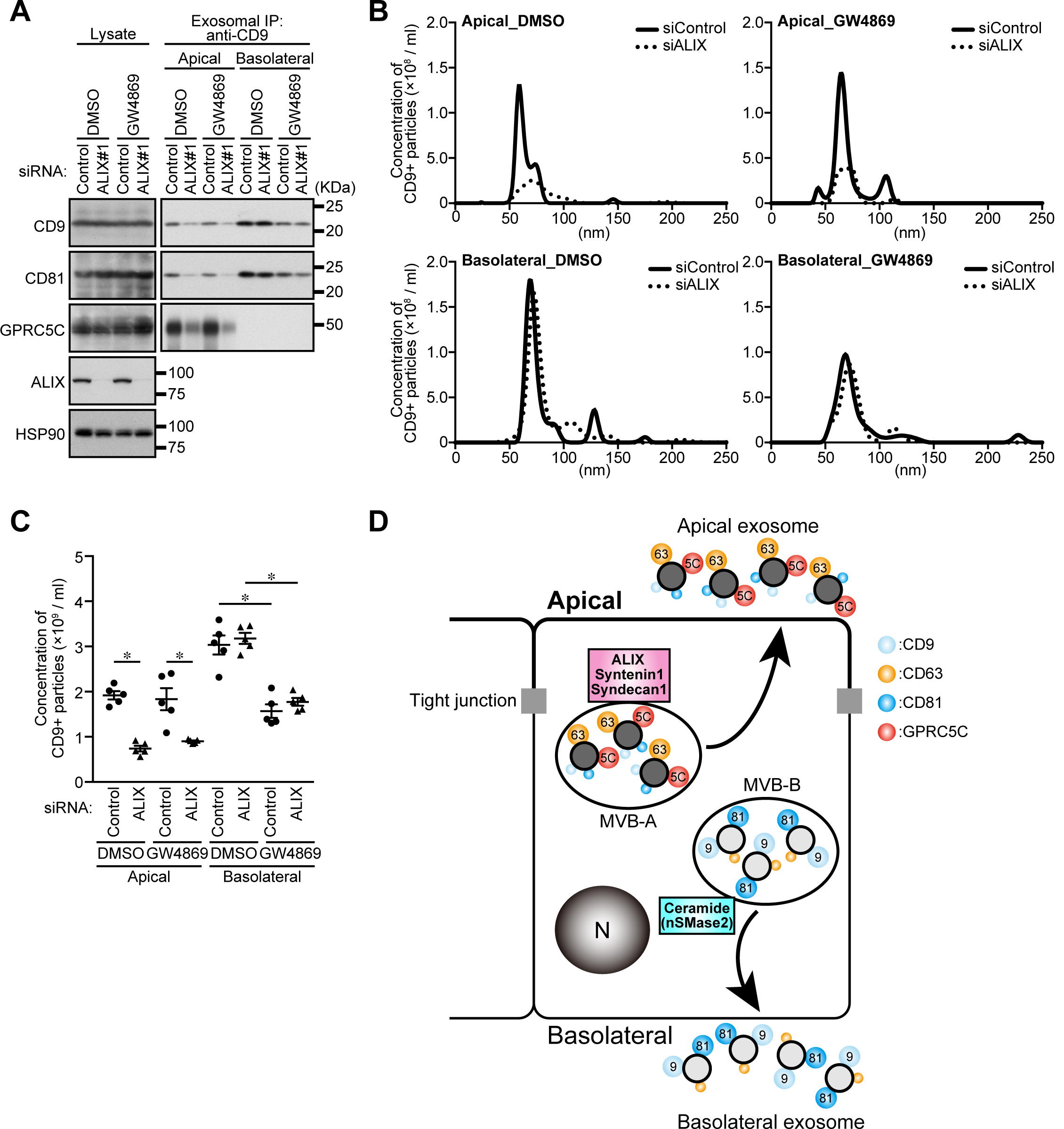
ALIX and ceramide independently mediate apical and basolateral exosome release, respectively. A. MDCK cells were transfected with siControl or siALIX, and the cells were transferred to cell culture inserts and cultured for 4 days. On the last day, the culture medium was replaced with EV-depleted medium with or without 10 nM GW4869. Exosomes were isolated from the pre-cleared medium by direct immunoaffinity capture using anti-CD9 antibody. Cell lysates and exosome samples were analyzed by immunoblotting with the antibodies indicated. B. Exosomes prepared as in (A) were eluted from the beads with a glycine buffer and analyzed by NTA. Representative NTA traces were shown. C. Quantification of the NTA data obtained in five independent experiments. \**P* < 0.01 (one-way ANOVA and Tukey’s test). Mean ± s.e.m. was shown. D. Model of polarized exosome release from MDCK cells. Apical exosome release and basolateral exosome release are mediated by ALIX–Syntenin1–Syndecan1 and ceramide, respectively. Marker sizes on exosomes (colored circles) reflect the abundance of each exosome protein.

The results of the present study provide evidence that distinct exosomes (apical and basolateral exosomes) are separately generated in polarized MDCK cells via two independent mechanisms. Considering that the significance of the existence of several exosome production mechanisms has never been elucidated thus far, this study is the first to demonstrate that two independent mechanisms, i.e., the ALIX–Syntenin1–Syndecan1 machinery (apical side) and the sphingomyelinase-dependent ceramide production machinery, are used for polarized exosome release from epithelial cells. Although it still remains unclear whether heterogeneous exosomes are formed in a single MVB or different MVBs, our findings suggest that the source of the exosomal heterogeneity in polarized epithelial cells is different MVBs rather than a single MVB, because CD9 was not overlapped well with CD63 (Fig EV2A) in polarized MDCK cells. Moreover, since CD63- and CD9-positive exosomes were also separately formed via an unknown mechanism in dendritic cells (Kowal *et al*., 2016), the two mechanisms can be used to generate exosomal heterogeneity even in non-polarized cells. Thus, our discovery provides an important clue to understanding of the generation of exosomal heterogeneity in many cell types in addition to epithelial cells.

## Materials and Methods

### Cell culture

MDCK II cells were cultured in Dulbecco’s modified Eagle’s medium (DMEM) (Fujifilm Wako Pure Chemical) supplemented with 10% fetal bovine serum (FBS), and 50 μg/ml penicillin and streptomycin in a 5% CO_2_ incubator. To purify extracellular vesicles (EVs) and exosomes, 1×10^6^ cells were plated on cell culture inserts (140640, Thermo Fisher scientific), and 4 h later unadherent cells were removed by medium change. The culture medium was changed every 24 h, and after 3 days the cells were washed twice with PBS and once with DMEM without FBS. The cells were then cultured for 24 h in DMEM containing 1% EV-depleted FBS. EV-depleted FBS was obtained by ultracentrifugation at 100,000 × *g* for 24 h and filtration through a 0.22-μm filter. For siRNA transfection, cells were plated on a culture dish, and one day later, the cells were transfected with siRNAs. After an additional 24 h, the cells were transferred to cell culture inserts, and two days later the cells were again transfected with siRNAs. For drug treatments, cells were cultured with 10 nM GW4869 (Sigma-Aldrich) for 24 h or with 100 nM bafilomycin A_1_ (Sigma-Aldrich) for the times indicated.

### Antibodies and reagents

All primary antibodies used in this study are listed in Table S1. Anti-dog GPRC5C rabbit polyclonal antibody was raised against its C-terminal region (AA 298-442). Horseradish peroxidase (HRP)-conjugated anti-mouse IgG goat polyclonal antibody (Southern Biotech), HRP-conjugated anti-rabbit IgG donkey antibody and HRP-conjugated anti-goat IgG donkey antibody (GE Healthcare), HRP-conjugated Protein-G (abcam), and Alexa Fluor 488/555-conjugated anti-goat/mouse IgG donkey polyclonal antibodies (Thermo Fisher Scientific) were used as secondary antibodies.

### RNAi

siRNA oligonucleotides were purchased from Nippon Gene. The target sequences used are listed in Table S1. Cells were transfected with the siRNA oligonucleotides by using Lipofectamine RNAiMAX (Thermo Fisher Scientific) according to the manufacturer’s instructions.

### Isolation of extracellular vesicles from cultured cells in culture inserts

The collected media were first subjected to a centrifugation step of 700 × *g* for 5 min to pellet and remove cells, and the supernatant was spun at 3,000 × *g* for 10 min to remove cell debris and apoptotic bodies. The 2nd supernatant was re-centrifuged at 10,000 × *g* for 30 min to remove heavy microvesicles, and any remaining large EVs were removed by passing the final supernatant through a 0.22-μm pore PES filter (Millipore). The obtained supernatant (pre-cleared medium) was then subjected to polyethylene glycol (PEG) precipitation as described previously (Rider *et al*., 2016) or ultracentrifugation to isolate the EVs. For PEG precipitation, the equal volume of 0.22-μm filtered 16% PEG-6000 solution (16% PEG-6000 and 1 M NaCl) added to the pre-cleared medium and the mixed samples were then refrigerated overnight. The following day, the samples were centrifuged at 4,000 × *g* for 1 h. The pellets obtained were suspended in 0.22-μm filtered PBS, and the suspensions were ultracentrifuged at 100,000 × *g* for 1 h in MLS-50 rotor (Beckman Coulter) to wash the samples (final sample; PEG pellet) (Fig EV1). To isolate EVs by ultracentrifugation, the pre-cleared medium was subjected to ultracentrifugation at 100,000 × *g* for 1 h. The pellet obtained was washed with 0.22-μm filtered PBS the same as after PEG precipitation (final sample; P100) (Fig EV1). For immunoblotting, the final samples were lysed with an SDS sample buffer without reducing agent, and for nanoparticle tracking assay (NTA) or negative staining, the final sample was suspended in 0.22-μm filtered PBS.

### Direct immunoaffinity capture of exosomes

Direct immunoaffinity capture was performed as described previously (Jeppesen *et al*., 2019) (Jeppesen *et al*., 2019). In brief, the pre-cleared medium was split into three portions. One portion was incubated for 16 h at 4°C with Dynabeads (Thermo Fisher Scientific) directly conjugated to anti-CD9 antibody with rotation, and the second portion with Dynabeads conjugated to normal mouse IgG (Santa Cruz). The third portion was subjected to ultracentrifugation and washing to obtain P100 (as described above). After incubation, the beads were washed twice with 0.22-μm filtered ice-cold 0.1% BSA-PBS, and washed once with 0.22-μm filtered PBS. Immediately following the final wash, the exosome-loaded beads were suspended in an SDS sample buffer without reducing agent. The beads were removed from the suspension with a magnet, and the clarified lysates were used for immunoblotting. For NTA, exosomes were eluted from the beads as follows: after the final wash, the exosome-loaded beads were incubated in 0.22-μm filtered 0.1 M glycine-HCl buffer (pH 3.0) for 5 min at room temperature and then equilibrated with one-fourth volume of 0.22-μm filtered 1 M Tris-HCl (pH 8.0). The supernatant that remained after removing of the beads with the magnet was suspended in 0.22-μm filtered PBS.

### Immunoblotting

Cells were collected with an SDS sample buffer without reducing agent and dispersed through a 25-gauge needle. Samples were subsequently separated by SDS-PAGE and transferred to Immobilon-P polyvinylidene difluoride membranes (Millipore). Immunoblot analysis was performed with the antibodies indicated and visualization was achieved with the Immobilon Western Chemiluminescent HRP substrate (Millipore).

### Nanoparticle tracking assay

Purified EV or exosome samples were analyzed for particle concentration and size distribution by using the NTA method by Malvern NanoSight NS300 (Malvern Panalytical). The assays were performed according to the protocol recommended by the manufacturer. Briefly, three independent replicates of exosome preparations diluted in PBS (see above) were injected at a constant rate into the tracking chamber with the syringe pump provided. The specimens were tracked at room temperature for 60 s. Shutter and gain were manually adjusted for optimal detection and kept at the optimized settings for all samples. The data were captured and analyzed with NTA software (version 3.4, Malvern Panalytical).

### Floatation assay

PEG pellets were suspended in 0.5 ml of an ice-cold homogenization buffer (250 mM sucrose, 20 mM HEPES-KOH [pH 7.4], 1 mM EDTA, and complete EDTA-free protease inhibitor). The suspensions were diluted with an equal volume of 50% OptiPrep (Cosmo Bio) in the homogenization buffer. Discontinuous OptiPrep gradients were generated in MLS-50 tubes (Beckman Coulter) by overlaying the following OptiPrep solutions in the homogenization buffer: 1 ml of the diluted sample in 25% OptiPrep, 1.5 ml in 20% OptiPrep, 1.875 ml in 10% OptiPrep, and 0.625 ml in 0% OptiPrep. The gradients were centrifuged at 150,000 × *g* in MLS-50 rotors for 3 h, and then 10 fractions (0.5 ml each) were collected from the top. Proteins in each fraction were isolated by TCA precipitation. The final pellets were suspended in an SDS sample buffer without reducing agent.

### Mass spectrometry and data analysis

Proteins in P100, which was obtained from culture medium without FBS to avoid serum protein contamination, were concentrated by TCA (trichloroacetic acid) precipitation. The precipitates were suspended in 50 mM TEAB and 0.1% SDS, and the protein concentrations were determined by the BCA method. A 2 μg sample of the proteins from each suspension was reduced by adding DTT to a final concentration of 134 mM and incubating at 35°C for 2 h. Free thiol groups were alkylated by adding 230 mM iodoacetic acid and allowing to stand at room temperature for 30 min in the dark. Samples were digested with trypsin (APRO Science) overnight at 37°C, and the digested samples were prepared for mass spectrometry analysis by passage through a GL-Tip SCX (GL Science). The samples were reconstituted in a starting buffer composed of 10 mM KH_2_PO_4_ (pH 3.0), 25% acetonitrile, and 10 mM KCl. The peptides were eluted off the elution buffer of 10 mM KH_2_PO_4_ (pH 3.0), 25% acetonitrile, and 350 mM KCl. Each eluate was concentrated by vacuum centrifugation and resuspended in 50 μL of 0.1% (v/v) formic acid, and the samples were desalted using SPE C-Tip (Nikkyo Technos). The desalted samples were concentrated by vacuum centrifugation and resuspended in 40 μL of 0.1% (v/v) formic acid. All samples were stored at −20°C until LC-MS analysis. The peptides recovered were analyzed with a Q Exactive Plus mass spectrometer (Thermo Fisher Scientific) coupled on-line with a capillary high-performance liquid chromatography (HPLC) system (EASY-nLC 1200, Thermo Fisher Scientific) to acquire MS/MS spectra. For electrospray ionization, a 0.075 × 150 mm-EASY-Spray column (3-μm particle diameter, 100 Å pore size, Thermo Fisher Scientific) with mobile phases of 0.1% formic acid and 0.1% formic acid/80% acetonitrile was used. Data derived from the MS/MS spectra were used to search the SWISS-Prot protein database by using the MASCOT Server [http://www.matrixscience.com] and to identify proteins by using the Scaffold viewer program [http://www.proteomesoftware.com/products/scaffold].

### Immunocytochemistry

Cells grown on coverslips were washed with PBS and fixed in 10% TCA for 10 min at - 30°C. The fixed cells were permeabilized with 50 μg/ml digitonin (Sigma-Aldrich) in PBS for 3 min, blocked with 3% bovine serum albumin in PBS for 30 min, and then incubated with anti-CD63 goat and/or anti-CD9 mouse antibodies for 1 h. After washing three times with PBS, the cells were incubated with Alexa Fluor 488-conjugated anti-goat IgG and Alexa Fluor 555-conjugated anti-mouse IgG secondary antibodies for 1 h. The coverslips were observed using a confocal laser microscope (FV1000 IX81, Olympus) with a 100 × oil-immersion objective lens (1.45 NA, Olympus) and captured with FluoView software (Olympus). The images were processed by using Photoshop 2020 software (Adobe).

### Electron microscopy

Cells were cultured on cell tight C-2 cell disks (Sumitomo Bakelite) and fixed for 2 h in 2.5% glutaraldehyde (Electron Microscopy Science) in 0.1 M phosphate buffer (pH 7.4) on ice. The cells were washed with 0.1 M phosphate buffer (pH 7.4) three times, postfixed in 1% osmium tetroxide in 0.1 M phosphate buffer (pH 7.4) for 2 h, dehydrated, and embedded in Epon 812 according to the standard procedure. Ultrathin sections were stained with uranyl acetate and lead citrate. For immunonegative staining, a PEG pellet suspended in PBS was added to a nickel-coated formvar grid for 5 min. The excess solution was soaked off with a filter paper, and the samples were washed with 0.1 M phosphate buffer (pH 7.4) three times, then postfixed in 1% PFA (TAAB Laboratories Equipment) in 0.1 M phosphate buffer (pH 7.4) for 3 min. The grids were transferred to a solution containing mouse anti-CD9 antibody diluted 1:2000 for 2 h and then rinsed with 0.1 M phosphate buffer (pH 7.4) six times. Bound antibodies were detected with goat anti-mouse IgG (H+L) 5-nm colloidal Gold particles (BBI Solutions) and rinsed with 0.1 M phosphate buffer (pH 7.4) six times. The grids were washed twice with distilled water and negatively stained with 1% uranyl acetate for 2 min. All samples were examined with an H-7100 electron microscope (Hitachi).

### Statistical analysis

Two groups of data were evaluated by the unpaired two-tailed Student’s *t*-test, and multiple comparisons were performed by one-way analysis of variance (ANOVA) followed by the Tukey’s test. Statistical analysis was performed with Prism6 (GraphPad software).

## Acknowledgments

We thank K. Shoji for technical assistance, and E. Morita, N. Tanaka, and all members of the laboratory for helpful discussions. This work was supported in part by Grant-in-Aid for Research Activity Start-up 19K21174 from the Ministry of Education, Culture, Sports, Science and Technology (MEXT) of Japan (to T. M.); the Kao Foundation for Arts and Sciences (to T. M.); Grant-in-Aid for Scientific Research (B) 19H03220 from MEXT (to M. F.); and Japan Science and Technology Agency (JST) CREST Grant JPMJCR17H4 (to M. F.).

## Author Contributions

T. M, and M. F. designed the experiments, interpreted the data, and wrote the manuscript. T. M, S. H, and F. O. carried out the experiments and interpreted the data. Y.S. performed EM analysis. All authors discussed the results and commented on the manuscript.

## Expanded view figure legends

**Figure EV1. Scheme of the EV and exosome isolation methods used in this study**.

**Figure EV2. Localization and structure of MVBs in polarized MDCK cells**.

A. Polarized MDCK cells were immunostained with anti-CD63 and anti-CD9 antibodies. Arrowheads, CD63- and CD9-double-positive dots; arrows, CD63-positive dots; and double-headed arrows, CD9-positive dots. Scale bar, 20 μm.

B. Polarized MDCK cells were analyzed by conventional electron microscopy. Small ILVs and a large ILV were indicated by arrowheads and an arrow, respectively. It is noteworthy that almost all of the ILVs measured <100 nm in diameter. Scale bar, 100 nm.

C. MDCK cells were cultured on cell culture inserts for 4 days. On the last day, the culture medium was replaced with EV-depleted medium. EVs released from the apical and basolateral side of the MDCK cells were purified by PEG precipitation and analyzed by NTA. Representative NTA traces were shown.

D. Quantification of the NTA data obtained in five independent experiments. Note that both PEG pellets showed a broad particle size distribution, suggesting that PEG pellets contain various types of EVs.

**Figure EV3. Exosome release is upregulated by inhibition of lysosomal function**

A. MDCK cells were cultured on cell culture inserts for 4 days. On the last day, the cells were cultured in EV-depleted medium with or without 100 nM bafilomycin A_1_ (Baf A_1_) for the times indicated until the medium was harvested. Exosomes were isolated from the pre-cleared medium by direct immunoaffinity capture using anti-CD9 antibody. Cell lysates and exosome samples were analyzed by immunoblotting with the antibodies indicated.

B. Exosomes prepared as in (A) were eluted from the beads with a glycine buffer and analyzed by NTA. Representative NTA traces were shown.

C. Quantification of the NTA data obtained in five independent experiments. \**P* < 0.01 (one-way ANOVA and Tukey’s test). Mean ± s.e.m. was shown.

D. Polarized MDCK cells were cultured in EV-depleted medium with or without 100 nM Baf A_1_ for 6 h and cells were immunostained with anti-CD63 antibody. Scale bars, 20 μm (1 μm, insets).

E. MDCK cells were transfected with siControl or the siRNAs indicated. After 3 days, the culture medium was replaced with EV-depleted medium. One day later, the cells were immunostained with anti-CD63 antibody. Scale bars, 20 μm (1 μm, insets).

F. MDCK cells were transfected with the siRNAs indicated, cultured as in (E), and analyzed by conventional electron microscopy. Scale bars, 400 nm. Note that in addition to normal MVBs, enlarged and ILV-less MVB-like structures were often observed in HRS-KD cells.

G. MDCK cells were transfected with siControl or two independent siALIX. The cells were then transferred to cell culture inserts and cultured for 4 days. On the last day, the culture medium was replaced with EV-depleted medium. Exosomes were isolated from the pre-cleared medium by direct immunoaffinity capture using anti-CD9 antibody. Cell lysates and exosome samples were analyzed by immunoblotting with the antibodies indicated.

**Figure EV4. Syntenin1 and Syndecan1 are specifically involved in the apical exosome release**.

A. MDCK cells were transfected with siControl or the siRNAs indicated. The cells were then transferred to cell culture inserts and cultured for 4 days. On the last day, the culture medium was replaced with EV-depleted medium. Exosomes were isolated from the pre-cleared medium by direct immunoaffinity capture using anti-CD9 antibody. Cell lysates and exosome samples were analyzed by immunoblotting with the antibodies indicated.

B. Exosomes prepared as in (A) were eluted from the beads with a glycine buffer and analyzed by NTA. Representative NTA traces were shown.

C. Quantification of the NTA of data obtained in five independent experiments. \**P* < 0.05, \*\**P* < 0.01 (one-way ANOVA and Tukey’s test). Mean ± s.e.m. was shown.

**Table EV1. Proteins detected only in apically released EVs. Table EV2. Proteins detected only in basolaterally released EVs**.

**Table EV3. Proteins detected in apically and basolaterally released EVs**.

**Appendix Table S1. A list of materials used in this study**.

