## Supplementary material for "ALIX and ceramide differentially control polarized exosome release from epithelial cells": Table EV1

**Table EV1. Proteins detected only in apically released EVs.**

| Ranking of the detected peptide number | Protein name |
|---|---|
| 1 | G-protein coupled receptor family C group 5 member C isoform X1 |
| 2 | Golgi apparatus protein 1 |
| 3 | dihydropyrimidinase-related protein 2 |
| 4 | podocalyxin-like protein 1 precursor |
| 5 | myoferlin isoform X3 |
| 6 | brain-specific angiogenesis inhibitor 1-associated protein 2 |
| 7 | mucin-1 precursor |
| 8 | mucin-5B isoform X2 |
| 9 | LOW QUALITY PROTEIN: dipeptidyl peptidase 4 |
| 10 | Niemann-Pick C1-like 1 protein |
| 11 | pre-mRNA-processing-splicing factor 8 |
| 12 | heterochromatin protein 1-binding protein 3 isoform X1 |
| 13 | LOW QUALITY PROTEIN: epidermal growth factor receptor kinase substrate 8 |
| 14 | transmembrane 9 superfamily member 3 |
| 15 | unconventional myosin-VI isoform X2 |
| 16 | polymeric immunoglobulin receptor precursor |
| 17 | receptor-type tyrosine-protein phosphatase eta isoform X1 |
| 18 | LUC7-like 2 |
| 19 | 10 kDa heat shock protein, mitochondrial |
| 20 | 26S proteasome non-ATPase regulatory subunit 11 |
| 21 | transmembrane channel-like protein 5 isoform X1 |
| 22 | Sec1 homolog |
| 23 | lupus La protein |
| 24 | V-type proton ATPase catalytic subunit A |
| 25 | polypeptide N-acetylgalactosaminyltransferase 2 |
| 26 | U5 small nuclear ribonucleoprotein 200 kDa helicase |
| 27 | carboxypeptidase D |
| 28 | cytosolic 10-formyltetrahydrofolate dehydrogenase |
| 29 | COP9 signalosome complex subunit 4 isoform X1 |
| 30 | 60 kDa heat shock protein, mitochondrial |
| 31 | 3'-phosphoadenosine 5'-phosphosulfate transporter 1 |
| 32 | choline transporter-like protein 4 |
| 33 | CD109 antigen isoform X1 |
| 34 | charged multivesicular body protein 2b |
| 35 | galectin-9 |
| 36 | calcium and integrin-binding protein 1 isoform X2 |
| 37 | calpain-7 |
| 38 | aspartate--tRNA ligase, cytoplasmic isoform X1 |
| 39 | 26S proteasome regulatory subunit 10B |
| 40 | proliferating cell nuclear antigen |
| 41 | coatomer subunit gamma-1 |
| 42 | cytochrome P450 4A37 |
| 43 | coatomer subunit beta' isoform X1 |
| 44 | isocitrate dehydrogenase [NADP] cytoplasmic |
| 45 | protein FAM3C |
| 46 | CAD protein isoform X1 |
| 47 | vesicle-trafficking protein SEC22b |
| 48 | ruvB-like 2 |
| 49 | endothelial cell-specific molecule 1 |
| 50 | coatomer subunit alpha, partial |
| 51 | multidrug resistance p-glycoprotein |
| 52 | eukaryotic translation initiation factor 2 subunit 3 |
| 53 | UDP-glucose 6-dehydrogenase |
| 54 | vacuolar protein sorting-associated protein 4A isoform X2 |
| 55 | alpha-mannosidase 2 |
| 56 | adenylosuccinate lyase isoform X1 |
| 57 | melanotransferrin |
| 58 | COP9 signalosome complex subunit 1, partial |
| 59 | oxysterol-binding protein-related protein 3 isoform X1 |
| 60 | exportin-7 isoform X1 |
| 61 | apoptosis-inducing factor 1, mitochondrial isoform X1 |
| 62 | signal recognition particle,72 kDa subunit |
| 63 | mitochondrial import receptor subunit TOM70 isoform X2 |
| 64 | 26S proteasome non-ATPase regulatory subunit 12 isoform X1 |
