## Supplementary material for "ALIX and ceramide differentially control polarized exosome release from epithelial cells": Table EV2

**Table EV2. Proteins detected only in basolaterally released EVs.**

| Ranking of the detected peptide number | Protein name |
|---|---|
| 1 | extracellular matrix protein FRAS1 |
| 2 | calsyntenin-1 isoform X2 |
| 3 | coagulation factor V |
| 4 | FERM, RhoGEF and pleckstrin domain-containing protein 1 isoform X1 |
| 5 | integrin alpha-1 |
| 6 | hepatocyte growth factor-regulated tyrosine kinase substrate isoform 1 |
| 7 | glutaredoxin-3 isoform X2 |
| 8 | tensin-1 isoform X5 |
| 9 | nidogen-1 |
| 10 | far upstream element-binding protein 2 |
| 11 | LOW QUALITY PROTEIN: suppressor of tumorigenicity 14 protein |
| 12 | utrophin |
| 13 | fibulin-7 isoform X1 |
| 14 | bone morphogenetic protein 1 |
| 15 | semaphorin-4D isoform X1 |
| 16 | cartilage oligomeric matrix protein |
| 17 | src substrate cortactin isoform X1 |
| 18 | 4F2 cell-surface antigen heavy chain |
| 19 | coagulation factor XIII A chain |
