## Supplementary material for "ALIX and ceramide differentially control polarized exosome release from epithelial cells": Table EV3

**Table EV3. Proteins detected in apically and basolaterally released EVs.**

| Ranking of the detected peptide number | Protein name |
|---|---|
| 1 | basement membrane-specific heparan sulfate proteoglycan core protein, partial |
| 2 | fibronectin isoform X4 |
| 3 | agrin isoform X1 |
| 4 | glycoprotein 80 |
| 5 | thrombospondin-1 |
| 6 | ubiquitin-40S ribosomal protein S27a |
| 7 | unnamed protein product (CAR80291.1) |
| 8 | laminin subunit gamma-1 |
| 9 | unnamed protein product, partial |
| 10 | programmed cell death 6-interacting protein isoform X2 |
| 11 | prostaglandin F2 receptor negative regulator, partial |
| 12 | latent-transforming growth factor beta-binding protein 1 isoform X1 |
| 13 | lamin |
| 14 | actin, cytoplasmic 2 |
| 15 | myosin, heavy polypeptide 9, non-muscle |
| 16 | LOW QUALITY PROTEIN: collagen alpha-1(XVIII) chain |
| 17 | pyruvate kinase PKM isoform X1 |
| 18 | soluble calcium-activated nucleotidase 1 isoform X1 |
| 19 | keratin, type II cytoskeletal 8 |
| 20 | laminin subunit beta-1 |
| 21 | heat shock cognate 71 kDa protein |
| 22 | disintegrin and metalloproteinase domain-containing protein 10 |
| 23 | gelatinase B |
| 24 | LOW QUALITY PROTEIN: alpha-enolase |
| 25 | LOW QUALITY PROTEIN: laminin subunit alpha-5 |
| 26 | protein CYR61 isoform X2 |
| 27 | glyceraldehyde-3-phosphate dehydrogenase |
| 28 | major vault protein |
| 29 | neuroblast differentiation-associated protein AHNAK isoform X1 |
| 30 | annexin 2 |
| 31 | Na, K-ATPase alpha-1 subunit |
| 32 | integrin beta-1 isoform X2 |
| 33 | alpha-2-HS-glycoprotein |
| 34 | phosphoglycerate kinase 1 |
| 35 | LOW QUALITY PROTEIN: lysyl oxidase homolog 2 |
| 36 | follistatin |
| 37 | filamin-A |
| 38 | alpha-2-macroglobulin |
| 39 | T-complex protein 1 subunit theta isoform X2 |
| 40 | keratin, type I cytoskeletal 19 |
| 41 | vimentin |
| 42 | neurofilament light polypeptide |
| 43 | neurofilament medium polypeptide |
| 44 | RecName: Full=Serum albumin; AltName: Allergen=Can f 3; Flags: Precursor |
| 45 | stanniocalcin-1 |
| 46 | Na(+)/H(+) exchange regulatory cofactor NHERF1 |
| 47 | histone H1.4 |
| 48 | transcobalamin-2 |
| 49 | syntenin-1 isoform X1 |
| 50 | transitional endoplasmic reticulum ATPase isoform X2 |
| 51 | annexin A1 |
| 52 | basal cell adhesion molecule |
| 53 | insulin-like growth factor-binding protein 7 |

| | |
|---|---|
| 54 | transgelin-2 |
| 55 | insulin-like growth factor-binding protein 4 |
| 56 | talin-1 isoform X2 |
| 57 | slit homolog 3 protein |
| 58 | vinculin |
| 59 | laminin subunit alpha-3 isoform X1 |
| 60 | ezrin |
| 61 | catenin alpha-1 |
| 62 | keratin, type II cytoskeletal 7 isoform X2 |
| 63 | C-1-tetrahydrofolate synthase, cytoplasmic isoform X2 |
| 64 | collagen alpha-2(V) chain isoform X1 |
| 65 | platelet glycoprotein IIIa |
| 66 | elongation factor 1-alpha 1 |
| 67 | retinal dehydrogenase 1 |
| 68 | T-complex protein 1 subunit delta |
| 69 | tubulin beta-4B chain isoform X2 |
| 70 | pregnancy zone protein-like isoform X1 |
| 71 | peroxiredoxin-1 |
| 72 | heterogeneous nuclear ribonucleoprotein K |
| 73 | fructose-bisphosphate aldolase A |
| 74 | integrin alpha-V |
| 75 | T-complex protein 1 subunit gamma isoform X1 |
| 76 | rab1 |
| 77 | membrane cofactor protein isoform X1 |
| 78 | epithelial cell adhesion molecule |
| 79 | podocan isoform X2 |
| 80 | fatty acid synthase |
| 81 | 14-3-3 protein zeta/delta |
| 82 | ras-related protein Rab-5B |
| 83 | elongation factor 2 |
| 84 | LOW QUALITY PROTEIN: T-complex protein 1 subunit epsilon |
| 85 | cofilin-1 |
| 86 | T-complex protein 1 subunit beta isoform X1 |
| 87 | beta-galactosides-binding lectin, partial |
| 88 | T-complex protein 1 subunit eta |
| 89 | IST1 homolog isoform X1 |
| 90 | L-lactate dehydrogenase A chain isoform X2 |
| 91 | LOW QUALITY PROTEIN: uncharacterized protein LOC488306 |
| 92 | versican core protein isoform X2 |
| 93 | keratin, type I cytoskeletal 18 |
| 94 | 40S ribosomal protein S2 isoform X1 |
| 95 | osteopontin, partial |
| 96 | annexin XIIIb |
| 97 | clathrin heavy chain 1 isoform X1 |
| 98 | lactadherin |
| 99 | probable ATP-dependent RNA helicase DDX17, partial (XP_022280011.1) |
| 100 | nucleobindin-1 |
| 101 | cyclophilin A, partial |
| 102 | laminin-5 gamma 2 |
| 103 | 14-3-3 protein sigma |
| 104 | core histone macro-H2A.1 isoform X2 |
| 105 | heat shock protein HSP 90-beta |
| 106 | macrophage-capping protein isoform X1 |
| 107 | vascular endothelial growth factor receptor 1 isoform X1 |
| 108 | laminin subunit alpha-5-like, partial |
| 109 | nucleolin isoform X1 |

| | |
|---|---|
| 110 | dnaJ homolog subfamily C member 3 |
| 111 | epithelial keratin 1 |
| 112 | a disintegrin and metallopeptidase domain 9 |
| 113 | prosaposin isoform X1 |
| 114 | ATP-citrate synthase isoform X1 |
| 115 | LOW QUALITY PROTEIN: ubiquitin-like modifier-activating enzyme 1 |
| 116 | uridine phosphorylase 1 |
| 117 | T-complex protein 1 subunit alpha isoform X2 |
| 118 | cytoplasmic dynein 1 heavy chain 1 |
| 119 | beta amyloid precursor protein isoform APP770 |
| 120 | triose-phosphate isomerase |
| 121 | integrin alpha-2 |
| 122 | proteasome subunit alpha type-1 |
| 123 | non-specific lipid-transfer protein isoform X2 |
| 124 | 14-3-3 protein epsilon isoform X2 |
| 125 | 40S ribosomal protein S11 |
| 126 | CD63 antigen |
| 127 | heterogeneous nuclear ribonucleoprotein L isoform X1 |
| 128 | 40S ribosomal protein S3a |
| 129 | T-complex protein 1 subunit zeta |
| 130 | antithrombin-III |
| 131 | 40S ribosomal protein S4, X isoform |
| 132 | heat shock protein HSP 90-alpha |
| 133 | folate receptor beta isoform X2 |
| 134 | charged multivesicular body protein 4b |
| 135 | echinoderm microtubule-associated protein-like 4 isoform X3 |
| 136 | heterogeneous nuclear ribonucleoprotein M isoform X4 |
| 137 | epithelial keratin 10 |
| 138 | erythrocyte band 7 integral membrane protein stomatin |
| 139 | laminin subunit beta-2 isoform X4 |
| 140 | 60S ribosomal protein L6 isoform X2 |
| 141 | gamma-glutamyltranspeptidase 1 |
| 142 | LOW QUALITY PROTEIN: heterogeneous nuclear ribonucleoprotein A1-like |
| 143 | tetraspanin-8 |
| 144 | 40S ribosomal protein S8 |
| 145 | alpha-fetoprotein |
| 146 | hsc70-interacting protein |
| 147 | 40S ribosomal protein S3 |
| 148 | moesin isoform X1 |
| 149 | LOW QUALITY PROTEIN: laminin subunit beta-3 |
| 150 | 60S ribosomal protein L5 isoform X1 |
| 151 | proteasome subunit alpha type-6 isoform X1 |
| 152 | periplakin isoform X1 |
| 153 | testis derived transcript |
| 154 | 14-3-3 protein eta |
| 155 | multifunctional protein ADE2 isoform X12 |
| 156 | multimerin-2 isoform X2 |
| 157 | heterogeneous nuclear ribonucleoproteins A2/B1 isoform X1 |
| 158 | heterogeneous nuclear ribonucleoprotein A3 isoform X1 |
| 159 | poly(rC)-binding protein 2 isoform X1 |
| 160 | syndecan-4 |
| 161 | annexin A4 |
| 162 | carbonic anhydrase 2 |
| 163 | A-kinase anchor protein 12 isoform X3 |
| 164 | transketolase |
| 165 | bifunctional glutamate/proline--tRNA ligase isoform X1 |

| | |
|---|---|
| 166 | LOW QUALITY PROTEIN: collagen alpha-2(IV) chain |
| 167 | LOW QUALITY PROTEIN: plexin-B2 |
| 168 | unnamed protein product, partial |
| 169 | nucleophosmin |
| 170 | 60S ribosomal protein L7a |
| 171 | heterogeneous nuclear ribonucleoproteins C1/C2 isoform X6 |
| 172 | lipolysis-stimulated lipoprotein receptor, partial |
| 173 | ladinin-1 isoform X2 |
| 174 | ras GTPase-activating-like protein IQGAP1 |
| 175 | 60S ribosomal protein L7 |
| 176 | calmodulin-3 isoform X1 |
| 177 | alpha-tubulin |
| 178 | DLA class I molecule |
| 179 | rac2 |
| 180 | complement C3 |
| 181 | junctional adhesion molecule A |
| 182 | neural cell adhesion molecule L1 isoform X1 |
| 183 | gelsolin |
| 184 | splicing factor, proline- and glutamine-rich |
| 185 | 14-3-3 protein theta |
| 186 | DNA topoisomerase 1 |
| 187 | proteasome subunit alpha type-5 |
| 188 | apolipoprotein B-100 |
| 189 | BRO1 domain-containing protein BROX |
| 190 | charged multivesicular body protein 2a isoform X1 |
| 191 | proteasome subunit beta type-6 |
| 192 | protein DJ-1 isoform X2 |
| 193 | 60S ribosomal protein L3 |
| 194 | met proto-oncogene precursor |
| 195 | heat shock protein 70 |
| 196 | 60S ribosomal protein L34 |
| 197 | complement factor I isoform X3 |
| 198 | glypican-1 |
| 199 | myosin light polypeptide 6 isoform X1 |
| 200 | 60S ribosomal protein L28 |
| 201 | cathepsin B isoform X1 |
| 202 | ras-related protein Rap-1b |
| 203 | 14-3-3 protein beta/alpha |
| 204 | ephrin type-B receptor 2 isoform X1 |
| 205 | plasminogen activator inhibitor 1 RNA-binding protein isoform X4 |
| 206 | cytoplasmic FMR1-interacting protein 2 isoform X1 |
| 207 | fibulin-1 isoform X1 |
| 208 | elongin-B |
| 209 | Golgi membrane protein 1 isoform X1 |
| 210 | basigin isoform X1 |
| 211 | tumor susceptibility gene 101 protein |
| 212 | serine/arginine-rich splicing factor 6 isoform X1 |
| 213 | vitamin D-binding protein |
| 214 | tubulointerstitial nephritis antigen-like |
| 215 | netrin-4 isoform X2 |
| 216 | dihydropyrimidinase-related protein 2 isoform X3 |
| 217 | cathepsin D |
| 218 | clathrin light chain A isoform X4 |
| 219 | gamma-synuclein |
| 220 | thyroid hormone receptor-associated protein 3 |
| 221 | elongation factor 1-gamma |

| | |
|---|---|
| 222 | LIM and SH3 domain protein 1 isoform X1 |
| 223 | chloride intracellular channel protein 4 |
| 224 | CDC42 GTP-binding protein |
| 225 | GDP dissociation inhibitor isoform 2 |
| 226 | beta-catenin |
| 227 | integrin alpha-6 isoform X1 |
| 228 | spectrin alpha chain, non-erythrocytic 1 isoform X2 |
| 229 | receptor of activated protein C kinase 1 |
| 230 | LOW QUALITY PROTEIN: integrin beta-4 |
| 231 | annexin A11 |
| 232 | solute carrier family 12 member 2, partial |
| 233 | 26S proteasome non-ATPase regulatory subunit 3 |
| 234 | small nuclear ribonucleoprotein Sm D2 |
| 235 | guanine nucleotide-binding protein G(k) subunit alpha |
| 236 | WD repeat domain 1 |
| 237 | ribosomal protein S18 |
| 238 | A disintegrin and metalloproteinase with thrombospondin motifs 1 |
| 239 | eukaryotic translation initiation factor 2 subunit 1 |
| 240 | 2',3'-cyclic-nucleotide 3'-phosphodiesterase isoform X1 |
| 241 | adenylyl cyclase-associated protein 1 isoform X1 |
| 242 | transforming growth factor-beta-induced protein ig-h3 |
| 243 | SAP domain-containing ribonucleoprotein |
| 244 | 26S proteasome regulatory subunit 7 |
| 245 | coronin-1C |
| 246 | unconventional myosin-Id isoform X1 |
| 247 | beta-type proteasome 7 subunit |
| 248 | heterogeneous nuclear ribonucleoprotein D0 isoform X4 |
| 249 | charged multivesicular body protein 1b |
| 250 | 40S ribosomal protein S9 |
| 251 | 60S ribosomal protein L35a |
| 252 | cadherin-1 precursor |
| 253 | nuclease-sensitive element-binding protein 1 |
| 254 | myosin-10 isoform X1 |
| 255 | AP-2 complex subunit beta isoform X2 |
| 256 | catenin delta-1 isoform X1 |
| 257 | granulins |
| 258 | staphylococcal nuclease domain-containing protein 1 |
| 259 | proteasome subunit beta type-4 |
| 260 | rab11 |
| 261 | choline transporter-like protein 2 isoform X2 |
| 262 | protein S100-A11 |
| 263 | 60S ribosomal protein L4 |
| 264 | N-Myc downstream regulated gene 1 |
| 265 | transforming protein RhoA precursor |
| 266 | splicing factor, partial |
| 267 | adhesion G-protein coupled receptor G1 |
| 268 | caveolin-1 |
| 269 | keratin, type II cytoskeletal 5 |
| 270 | eukaryotic initiation factor 4A-I |
| 271 | transaldolase |
| 272 | F-actin-capping protein subunit beta isoform X2 |
| 273 | far upstream element-binding protein 1 isoform X11 |
| 274 | spectrin beta chain, non-erythrocytic 1 isoform X2 |
| 275 | ribonuclease inhibitor |
| 276 | semaphorin-3F isoform X2 |
| 277 | protocadherin Fat 1 isoform X2 |

| | |
|---|---|
| 278 | collagen alpha-1(V) chain |
| 279 | 78 kDa glucose-regulated protein |
| 280 | peptidyl-prolyl cis-trans isomerase FKBP4 |
| 281 | laminin subunit alpha-5-like |
| 282 | serine/arginine-rich splicing factor 1 |
| 283 | heterogeneous nuclear ribonucleoprotein U-like protein 2 |
| 284 | stimulatory GTP binding protein |
| 285 | cytoplasmic aconitate hydratase |
| 286 | syntaxin-7 isoform X2 |
| 287 | 60S ribosomal protein L13 isoform X2 |
| 288 | Raf kinase inhibitor protein |
| 289 | complement C1s subcomponent |
| 290 | VIP36 (vesicular integral-membrane protein) |
| 291 | NKG2D ligand 1 |
| 292 | 60S ribosomal protein L13a |
| 293 | heterogeneous nuclear ribonucleoprotein U isoform X1 |
| 294 | monocarboxylate transporter 1 |
| 295 | coiled-coil domain-containing protein 80 |
| 296 | semaphorin-3E, partial |
| 297 | prolow-density lipoprotein receptor-related protein 1 isoform X1 |
| 298 | alpha-actinin-4 isoform X3 |
| 299 | interleukin enhancer-binding factor 3 isoform X4 |
| 300 | follistatin-related protein 1 |
| 301 | ribosomal protein L18, partial |
| 302 | tropomyosin alpha-4 chain isoform X3 |
| 303 | proprotein convertase subtilisin/kexin type 5 |
| 304 | serrate RNA effector molecule homolog isoform X4 |
| 305 | proteasome subunit beta type-5 |
| 306 | tropomyosin alpha-3 chain isoform X6 |
| 307 | inter-alpha-trypsin inhibitor heavy chain H2 |
| 308 | 40S ribosomal protein S12 |
| 309 | proteasome alpha subunit type 7, partial |
| 310 | elongation factor 1-beta |
| 311 | annexin A8 |
| 312 | GRP94 |
| 313 | filamin-B isoform X1 |
| 314 | integrin alpha-3 |
| 315 | unconventional myosin-Ib isoform X1 |
| 316 | dnaJ homolog subfamily A member 2 |
| 317 | 14-3-3 protein gamma |
| 318 | protein disulfide-isomerase A3 |
| 319 | keratin 14 |
| 320 | MARCKS-related protein |
| 321 | 40S ribosomal protein S6 |
| 322 | dystroglycan |
| 323 | serotransferrin |
| 324 | angiopoietin-related protein 4, partial |
| 325 | radixin isoform X2 |
| 326 | proteasome subunit beta type-1 |
| 327 | cadherin-3 |
| 328 | EH domain-containing protein 1 isoform X2 |
| 329 | ADP-ribosylation factor 1 |
| 330 | stress-induced-phosphoprotein 1 |
| 331 | pleckstrin homology-like domain family B member 1 isoform X15 |
| 332 | sodium bicarbonate cotransporter 3 isoform X3 |
| 333 | semaphorin-4B |

| | |
|---|---|
| 334 | fibulin-5 isoform X1 |
| 335 | inactive tyrosine-protein kinase 7 |
| 336 | phosphatidylinositol-binding clathrin assembly protein isoform X1 |
| 337 | synaptosomal-associated protein 23 |
| 338 | vacuolar protein sorting-associated protein 4B |
| 339 | actin-related protein 3 |
| 340 | Ribosomal protein S14, partial |
| 341 | ELAV-like protein 1 |
| 342 | thioredoxin-like isoform X1 |
| 343 | immunoglobulin superfamily member 8 isoform X2 |
| 344 | cysteine-rich motor neuron 1 protein isoform X2 |
| 345 | intercellular adhesion molecule-1, partial |
| 346 | carcinoembryonic antigen-related cell adhesion molecule 1 isoform 4L |
| 347 | Golgi-associated plant pathogenesis-related protein 1 isoform X1 |
| 348 | glycine--tRNA ligase isoform X1 |
| 349 | 26S proteasome non-ATPase regulatory subunit 13 |
| 350 | protein-lysine 6-oxidase isoform X1 |
| 351 | CE1 |
| 352 | polypyrimidine tract-binding protein 1 isoform X3 |
| 353 | eukaryotic translation initiation factor 3 subunit A isoform X2 |
| 354 | activated RNA polymerase II transcriptional coactivator p15 |
| 355 | preproendothelin-1 |
| 356 | fibrinogen gamma chain isoform X1 |
| 357 | GTP-binding protein (rab7) |
| 358 | EH domain-containing protein 4 |
| 359 | RNA-binding motif protein, X chromosome |
| 360 | adenosylhomocysteinase |
| 361 | 26S proteasome non-ATPase regulatory subunit 2 |
| 362 | 40S ribosomal protein SA |
| 363 | 40S ribosomal protein S19 |
| 364 | complement C1r subcomponent |
| 365 | polyadenylate-binding protein 1 isoform X1 |
| 366 | niban-like protein 1 |
| 367 | dnaJ homolog subfamily A member 1 |
| 368 | unconventional myosin-Ie |
| 369 | ras-related protein R-Ras2 isoform X2 |
| 370 | collagen alpha-1(IV) chain |
| 371 | septin-9 |
| 372 | arginine--tRNA ligase, cytoplasmic |
| 373 | calpain-2 catalytic subunit |
| 374 | transmembrane protease serine 11E |
| 375 | cathepsin Z |
| 376 | actin-related protein 2/3 complex subunit 1B |
| 377 | D-3-phosphoglycerate dehydrogenase |
| 378 | carbonyl reductase [NADPH] 1 |
| 379 | L-lactate dehydrogenase B chain |
| 380 | serine/threonine-protein phosphatase 2A 65 kDa regulatory subunit A alpha isoform isoform X1 |
| 381 | charged multivesicular body protein 1a |
| 382 | AP-2 complex subunit alpha-2 isoform X1 |
| 383 | coxsackievirus and adenovirus receptor transcriptional variant 4 |
| 384 | matrin-3 |
| 385 | myosin-14 isoform X1 |
| 386 | LOW QUALITY PROTEIN: desmoplakin |
| 387 | ribosome receptor |
| 388 | CMP-N-acetylneuraminate-beta-galactosamide-alpha-2,3-sialyltransferase 4 |
| 389 | eukaryotic initiation factor 4A-III |

| | |
|---|---|
| 390 | proliferation-associated protein 2G4 isoform X1 |
| 391 | 60S acidic ribosomal protein P0 |
| 392 | clathrin interactor 1 isoform X2 |
| 393 | histone H1.0 |
| 394 | zyxin |
| 395 | 26S proteasome regulatory subunit 8 |
| 396 | bone marrow stromal antigen 2 |
| 397 | eukaryotic translation initiation factor 2 subunit 2 isoform X1 |
| 398 | proteasome subunit beta type-2 isoform X1 |
| 399 | U1 small nuclear ribonucleoprotein 70 kDa isoform X1 |
| 400 | ubiquitin carboxyl-terminal hydrolase 5 isoform X3 |
| 401 | eukaryotic translation initiation factor 3 subunit D |
| 402 | disks large homolog 1 isoform X9 |
| 403 | UTP--glucose-1-phosphate uridylyltransferase isoform X1 |
| 404 | tight junction protein |
| 405 | CD44 antigen precursor |
| 406 | follistatin-related protein 3 isoform X2 |
| 407 | cell cycle and apoptosis regulator protein 2 isoform X3 |
| 408 | splicing factor 3B subunit 2 isoform X3 |
| 409 | septin-7 isoform X2 |
| 410 | serine/arginine-rich splicing factor 2 |
| 411 | vacuolar protein sorting-associated protein 37D |
| 412 | transcription elongation factor A protein 1 isoform X1 |
| 413 | ZO-1 MDCK |
| 414 | CD59 glycoprotein |
| 415 | vesicle-associated membrane protein-associated protein B/C isoform X1 |
| 416 | LOW QUALITY PROTEIN: RNA-binding protein FUS |
| 417 | protein kinase C and casein kinase substrate in neurons protein 2 isoform X6 |
| 418 | syntaxin-binding protein 3 isoform X1 |
| 419 | 116 kDa U5 small nuclear ribonucleoprotein component |
| 420 | peroxiredoxin-2 isoform X2 |
| 421 | aminoacyl tRNA synthase complex-interacting multifunctional protein 1 isoform X2 |
| 422 | serine/arginine-rich splicing factor 9 |
| 423 | cation-independent mannose-6-phosphate/insulin-like growth factor 2 receptor protein precursor |
| 424 | plasminogen activator inhibitor 1 precursor |
| 425 | non-POU domain-containing octamer-binding protein |
| 426 | glia-derived nexin |
| 427 | U2 small nuclear ribonucleoprotein B" |
| 428 | tropomyosin alpha-1 chain isoform X19 |
| 429 | myb-binding protein 1A isoform X1 |
| 430 | serine/arginine-rich splicing factor 5 isoform X1 |
| 431 | cullin-associated NEDD8-dissociated protein 1 |
| 432 | nephronectin isoform X1 |
| 433 | C-terminal-binding protein 2 isoform X2 |
| 434 | epithelial keratin 2e |
| 435 | U1 small nuclear ribonucleoprotein A |
| 436 | alpha-actinin-1 isoform X3 |
| 437 | tumor necrosis factor receptor superfamily member 6B |
| 438 | EH domain-containing protein 3 |
| 439 | v-src sarcoma Schmidt-Ruppin A-2 viral oncogene-like protein, partial |
| 440 | keratin, type II cytoskeletal 3 |
