## Appendix table S1 for "ALIX and ceramide differentially control polarized exosome release from epithelial cells"

**Appendix Table S1. A list of materials used in this study.**

| Antibodies | | |
|---|---|---|
| **Target name** | **Catalog No.** | **Company** |
| Flotillin-1 | 610820 | BD Bioscience |
| CD63 | AB0047-200 | SICGEN |
| CD9 | MA1-80307 | Thermo Fisher Scientific |
| CD81 | 555675 | BD Bioscience |
| TOMM20 | sc-136211 | Santa Cruz Biotechnology |
| Annexin I | 610066 | BD Bioscience |
| GPRC5C | | generated in this study |
| HRS | 10390-1-AP | Proteintech |
| TSG101 | sc-136111 | Santa Cruz Biotechnology |
| EAP20 (VPS25) | sc-271648 | Santa Cruz Biotechnology |
| EAP30 (VPS22) | sc-390747 | Santa Cruz Biotechnology |
| CHMP6 | 16278-1-AP | Proteintech |
| VPS4 | SAB4200025 | Sigma-Aldrich |
| ALIX | sc-53540 | Santa Cruz Biotechnology |
| $\beta$-actin | G043 | Applied Biological Materials |
| HSP90 | 610419 | BD Bioscience |
| Syntenin1 | PA5-42592 | Thermo Fisher Scientific |
| Syndecan1 | orb10289 | biorbyt |
| nSMase2 | ab85017 | Abcam |

| siRNAs | | |
|---|---|---|
| **siRNA name** | **Target gene** | **Target sequence** |
| siControl | Lusiferase | CGUACGCGGAAUACUUCGA |
| siALIX#1 | ALIX | AGAGCUGUGUGUUGUUCAA |
| siALIX#2 | ALIX | UCAGAUUUGUUCCAUUGAA |
| siCHMP6 | CHMP6 | GGAGCUGAAUGCAAUUACU |
| siEAP20 | EAP20 | AGCUCCCUGUGGAAUCAAU |
| siEAP30 | EAP30 | AGACCAACCUGGAGGAAUU |
| siHRS | HRS | AGAGGCAGGUGGAAGUAAA |
| sinSMase2 | nSMase2 | ACAGCAAGUUCCCCUACAA |
| siSyndecan1#1 | Syndecan1 | GAUCUCGGUUCUUGCAGAA |
| siSyndecan1#2 | Syndecan1 | CUUCACCUUUGACGUAUCU |
| siSyntenin1#1 | Syntenin1 | GCAAGACCUUCCAGUAUGA |
| siSyntenin1#2 | Syntenin1 | UAACAUCUGUGAAGUCAAU |
| siTSG101 | TSG101 | CAAUUCCUGUGCCUUAUAA |
| siVPS4A | VPS4A | UGGGUGCUGUUGUGAUAGA |
| siVPS4B | VPS4B | AGACUGAGUUCCUAGUUCA |
