## Supplementary figures and images for "ALIX and ceramide differentially control polarized exosome release from epithelial cells"

### Supplementary Figures

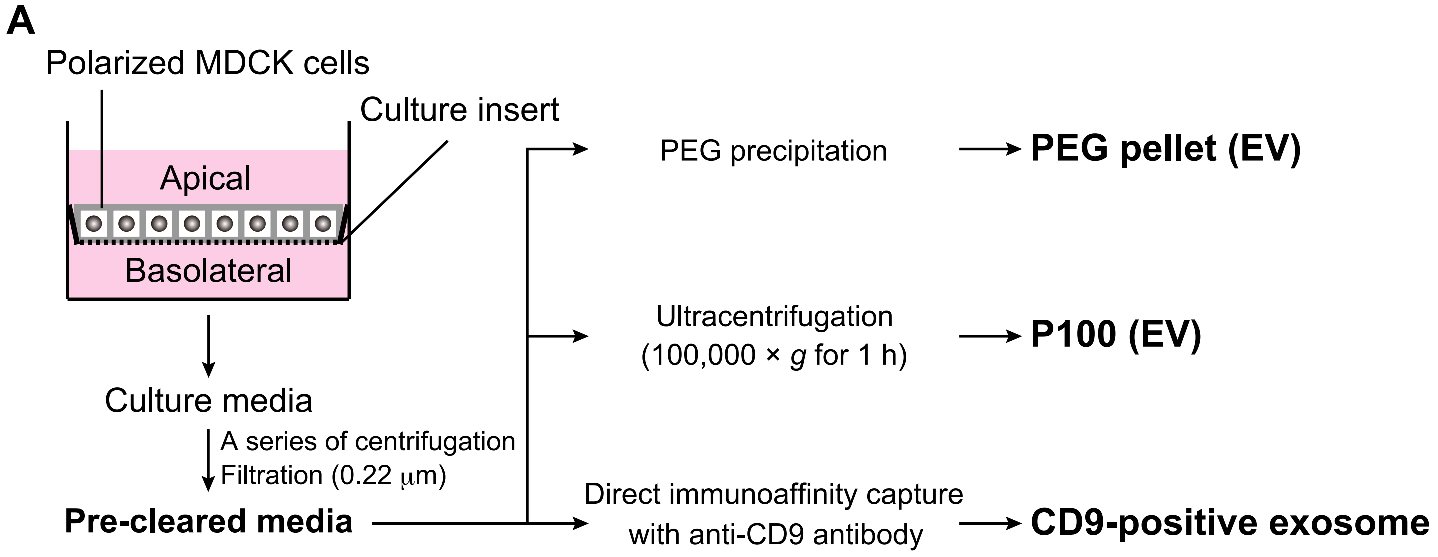

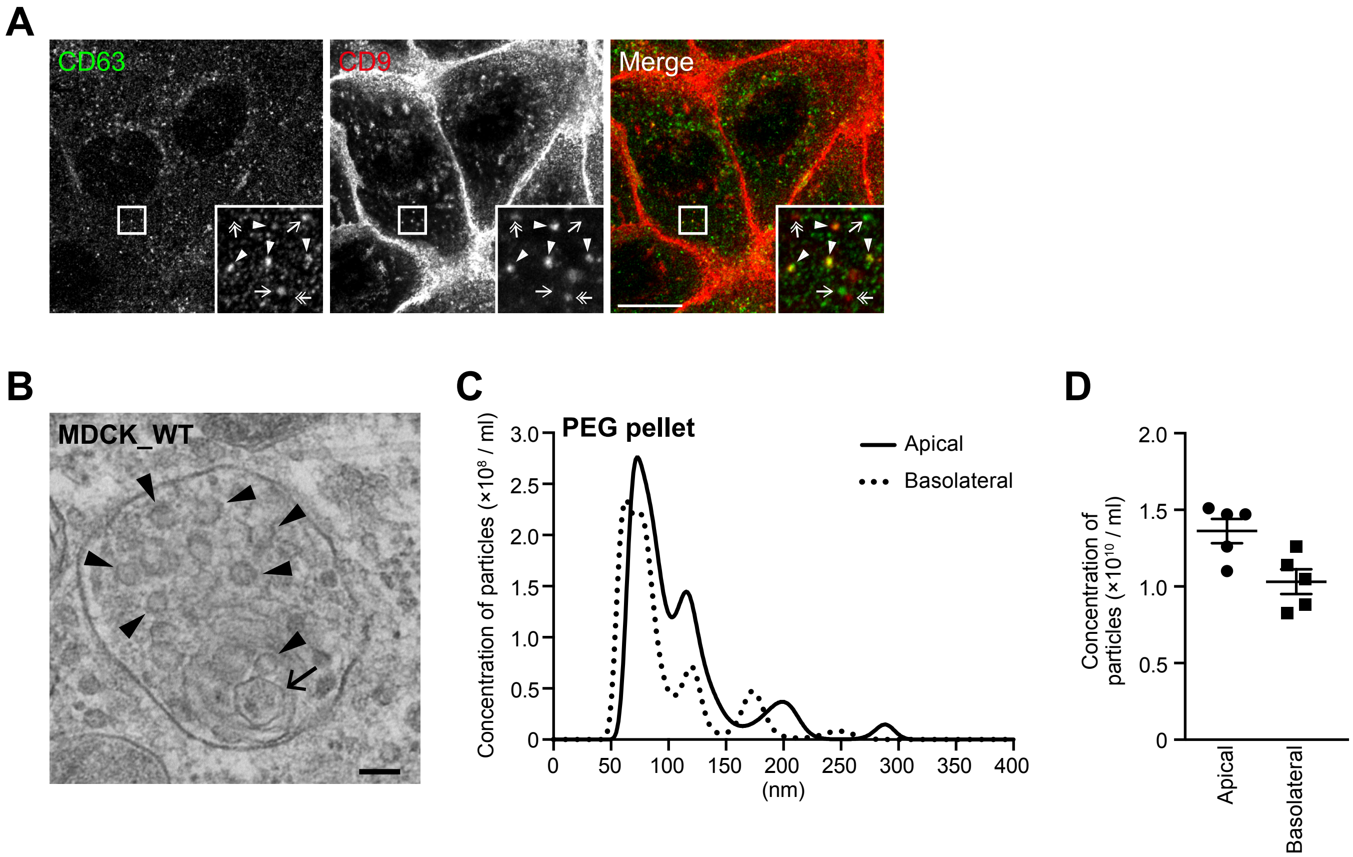

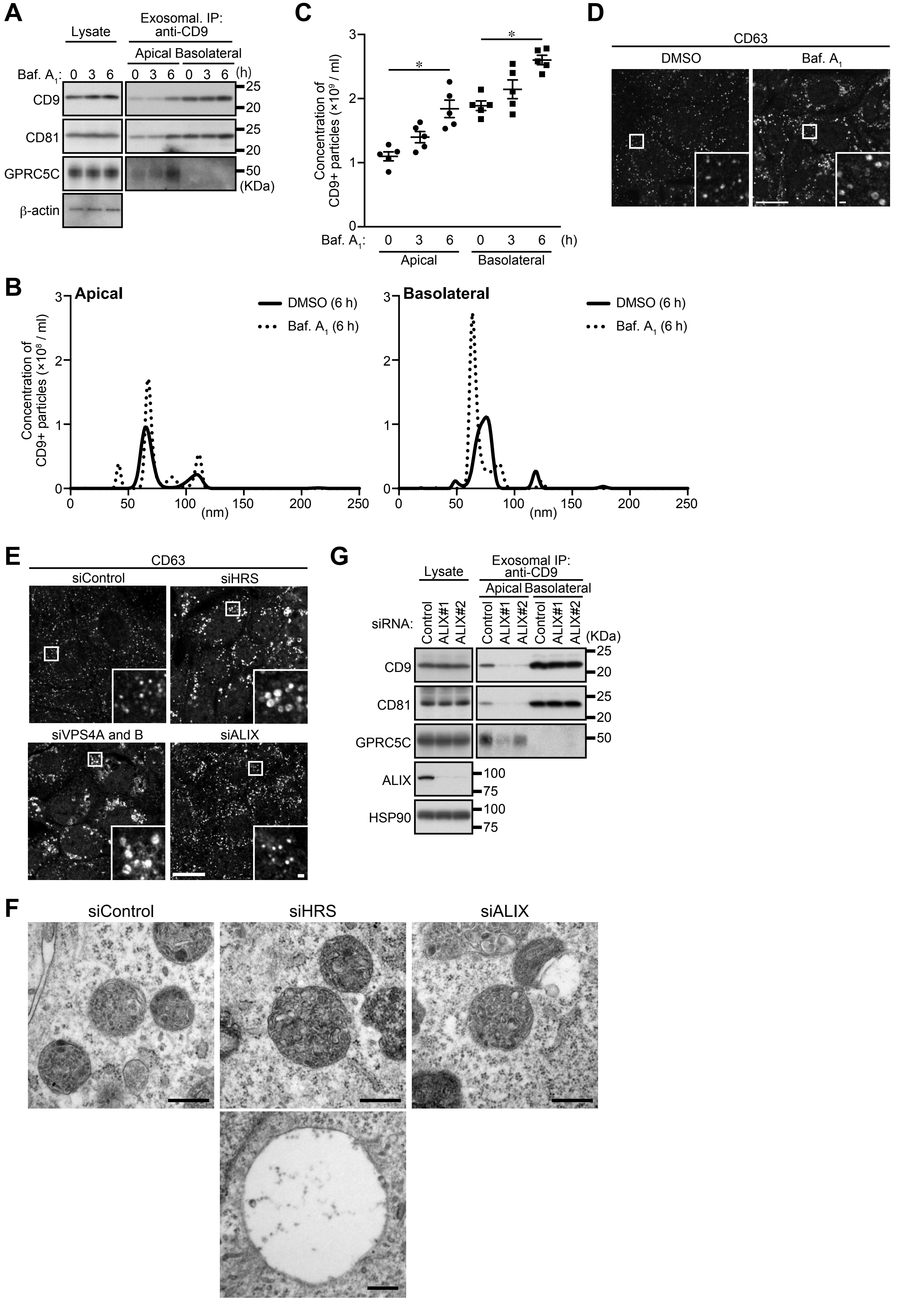

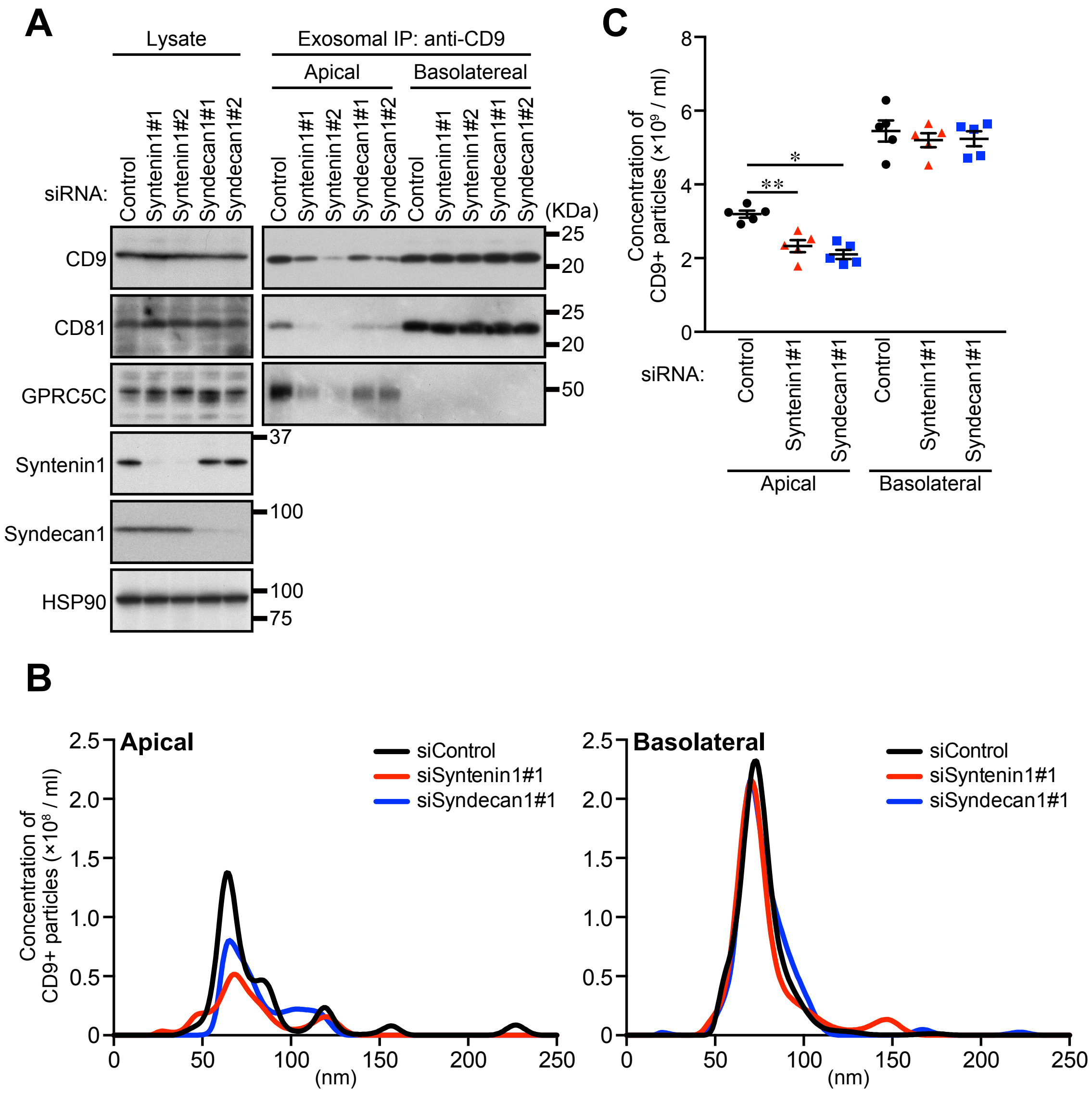

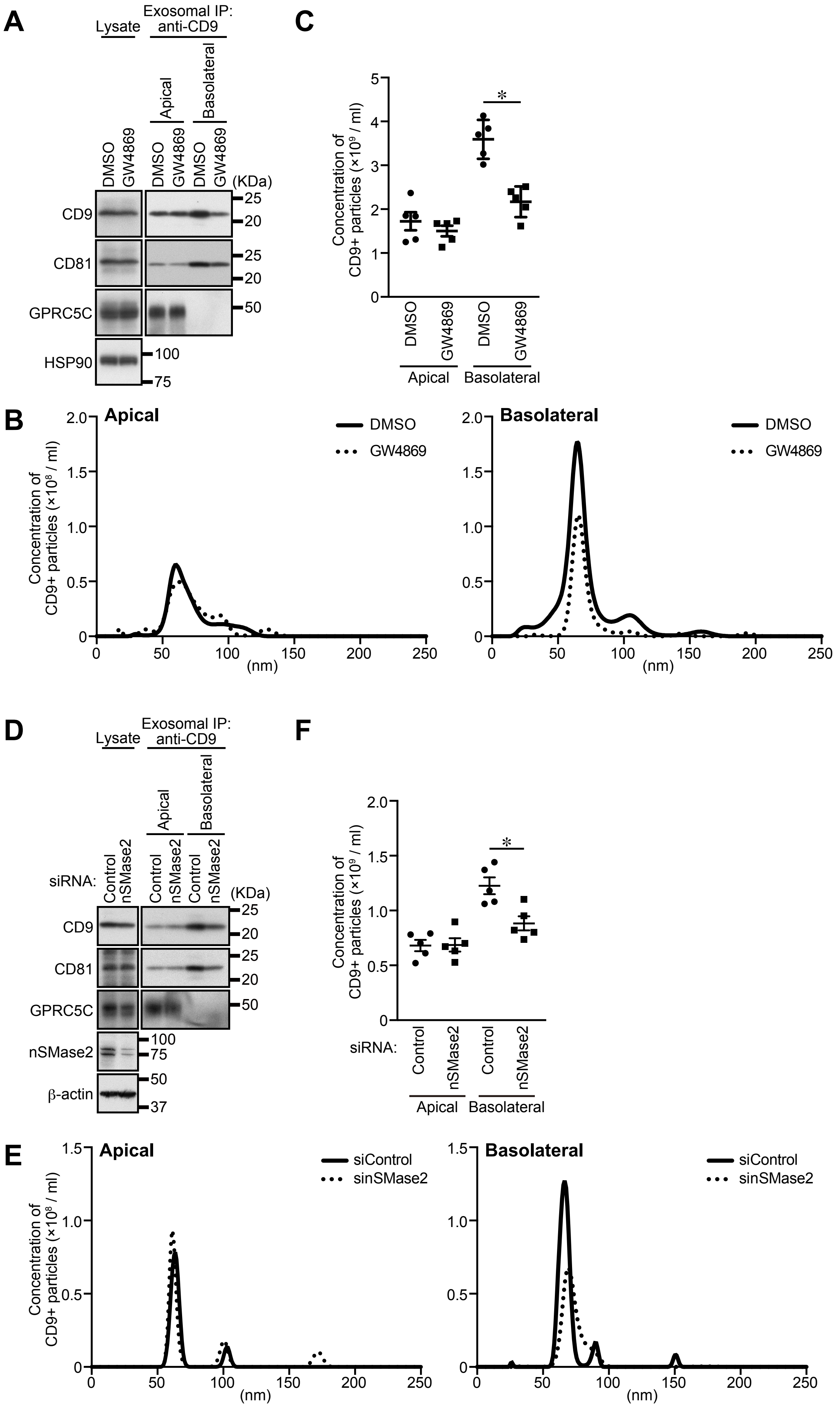
